# Neuropeptide Y expression defines a novel class of GABAergic projection neuron in the inferior colliculus

**DOI:** 10.1101/812388

**Authors:** Marina A. Silveira, Justin D. Anair, Nichole L. Beebe, Pooyan Mirjalili, Brett R. Schofield, Michael T. Roberts

## Abstract

Located in the midbrain, the inferior colliculus (IC) integrates information from numerous auditory nuclei and is an important hub for sound processing. Despite its importance, little is known about the molecular identity and functional roles of defined neuron types in the IC. Using a multifaceted approach in mice, we found that neuropeptide Y (NPY) expression identifies a major class of inhibitory neurons, accounting for approximately one-third of GABAergic neurons in the IC. Retrograde tracing showed that NPY neurons are principal neurons that can project to the medial geniculate nucleus. In brain slice recordings, many NPY neurons fired spontaneously, suggesting that NPY neurons may drive tonic inhibition onto postsynaptic targets. Morphological reconstructions showed that NPY neurons are stellate cells, and the dendrites of NPY neurons in the tonotopically-organized central nucleus of the IC cross isofrequency laminae. Immunostaining confirmed that NPY neurons express NPY, and we therefore hypothesized that NPY signaling regulates activity in the IC. In crosses between Npy1r^cre^ and Ai14 Cre-reporter mice, we found that NPY Y_1_ receptor (Y_1_R)-expressing neurons are glutamatergic and were broadly distributed throughout the rostro-caudal extent of the IC. In whole-cell recordings, application of a high affinity Y_1_R agonist led to hyperpolarization in most Y_1_R-expressing IC neurons. Thus, NPY neurons represent a novel class of inhibitory principal neurons that are well poised to use GABAergic and NPY signaling to regulate the excitability of circuits in the IC and auditory thalamus.

## Introduction

The inferior colliculus (IC) is the hub of the central auditory pathway (Adams, 1979; Cant and Benson, 2006, 2007), a critical processing center for most aspects of hearing, and an important site of plasticity after hearing loss (Chambers et al., 2016; Sturm et al., 2017). Despite its importance in auditory processing, it has proven difficult to identify the neuron classes that comprise the IC (Peruzzi et al., 2000; Palmer et al., 2013; Beebe et al., 2016). As a result, the cellular organization and function of neural circuits in the IC remain largely unclear.

The IC contains three main subdivisions: the central nucleus (ICc), dorsal cortex (ICd), and lateral cortex (IClc) (Morest and Oliver, 1984; Faye-Lund and Osen, 1985). The ICc is tonotopically organized into isofrequency laminae, and neurons in the ICc are divided in two broad morphological families: disc-shaped and stellate neurons (Meininger et al., 1986; Malmierca et al., 1993). Disc-shaped neurons, the majority of neurons, maintain their dendritic arbors within isofrequency laminae (Oliver and Morest, 1984), while stellate neurons extend their dendritic arbors across isofrequency laminae and integrate information across sound frequencies (Oliver et al., 1991). However, there is considerable diversity within the disc-shaped and stellate groups, suggesting that they contain multiple neuron types (Oliver et al., 1994; Peruzzi et al., 2000; Ono et al., 2005).

GABAergic neurons represent ∼25% of neurons in the IC (Oliver et al., 1994; Merchán et al., 2005; Beebe et al., 2016). In addition to providing local inhibition, many GABAergic neurons project to the medial geniculate nucleus (MG, auditory thalamus) (Winer et al., 1996; Peruzzi et al., 1997). Anatomical studies have subdivided GABAergic neurons into classes based on soma size and extracellular markers (Ito et al., 2009; Beebe et al., 2016), and neurons in all these classes can project to the MG (Beebe et al., 2018). However, it is unclear whether these anatomically defined groups represent neuron classes with distinct functional roles.

In many brain regions, an approach that combines anatomical and physiological analyses with molecular markers has proven key to identifying functionally distinct neuron types (Tremblay et al., 2016; Zeng and Sanes, 2017). Using such an approach, we recently identified vasoactive intestinal peptide (VIP) as a marker for a distinct class of excitatory principal neurons in the IC (Goyer et al., 2019), but there are currently no known molecular markers for GABAergic neuron classes in the IC.

NPY is a 36 amino-acid neuropeptide and one of the most abundant peptides in the brain (Chronwall et al., 1985; O’Donohue et al., 1985). NPY is released from neurons and binds to the NPY family of G_i/o_-coupled G-protein coupled receptors (Dumont et al., 1998). NPY signaling modulates synaptic transmission, neuronal excitability, and other physiological processes (Bacci et al., 2002). Across several brain regions, NPY has proven to be an important molecular marker for distinct classes of GABAergic neurons (Tong et al., 2008; Polgár et al., 2011). Because previous studies suggest that NPY (Nakagawa et al., 1995) and NPY Y_1_ receptors (Y_1_R) are expressed in the IC (Kishi et al., 2005; Eva et al., 2006), we hypothesized that NPY is a molecular marker for a class of GABAergic neurons in the IC.

Using a multifaceted approach, we found that NPY is expressed by a distinct population of GABAergic neurons in the IC. These neurons have a stellate morphology and represent approximately one-third of the GABAergic neurons in the IC. Retrograde tracing showed that NPY neurons can project to the MG. In brain slice recordings, NPY neurons fired spontaneously, suggesting a tonic release of NPY and GABA. Using Npy1r^cre^ mice, we found that Y_1_R-expressing IC neurons are glutamatergic and that application of a Y_1_R-selective agonist hyperpolarized the membrane potential of most Y_1_R-expressing neurons. Thus, our data indicate that NPY neurons represent a novel class of GABAergic IC principal neurons that may use GABAergic and NPY signaling to regulate the excitability of postsynaptic targets.

## Materials and Methods

### Animals

All experiments were approved by the University of Michigan Institutional Animal Care and Use Committee and were in accordance with NIH guidelines for the care and use of laboratory animals. Animals were kept on a 12-hour day/ night cycle with ad libitum access to food and water. NPY-hrGFP mice were obtained from Jackson Laboratory (stock # 006417; van den Pol et al., 2009) and were maintained hemizygous for the hrGFP transgene by crossing with C57BL/6J mice (Jackson Laboratory, stock # 000664). To visualize neurons that express Y_1_R, Npy1r^cre^ mice (B6.Cg-*Npy1r*^*tm1*.*1(cre/GFP)Rpa*^/ Jackson Laboratory, stock # 030544; Padilla et al., 2016) were crossed with Ai14 Cre-reporter mice (B6.Cg-*Gt(ROSA)26Sor* ^*tm14(CAG-tdTomato)Hze*^, Jackson Laboratory, stock # 007914). All mouse lines were on a C57BL/6J background. Because C57BL/6J mice are subject to age-related hearing loss due to the *Cdh23*^*ahl*^ mutation (Noben-Trauth et al., 2003), experiments were performed in animals aged P22 – P74, before the onset of hearing loss symptoms at about 3 months of age.

### Analysis of the distribution of NPY neurons

Three NPY-hrGFP mice were used to assess the distribution of NPY neurons in the IC. Each hrGFP-expressing IC neuron was identified in a series of every third section through the rostro-caudal extent of one IC. The locations of the labeled neurons were plotted using a Neurolucida system (MBF Bioscience) attached to a Zeiss Axio Imager.Z2 microscope. IC subdivisions were determined by examining staining for GAD67 and GlyT2 (Buentello et al., 2015). Low magnification photomicrographs of the distribution of NPY neurons were collected using a 5x objective on a Zeiss Axio Imager.Z2 microscope. Neurolucida Explorer was used to export data to Excel for analysis.

### Immunohistochemistry

Mice were deeply anesthetized and perfused transcardially with 0.1 M phosphate-buffered saline (PBS), pH 7.4, for 1 min and then with a 10% buffered formalin solution (Millipore Sigma, cat# HT501128) for 15 min. Brains were collected and post-fixed in the same fixative for 2 hours and cryoprotected overnight at 4°C in 0.1 M PBS containing 20% sucrose. Brains were cut into 40 μm sections on a vibratome or freezing microtome. Sections were washed in 0.1 M PBS, and then treated with 10% normal donkey serum (Jackson ImmunoResearch Laboratories, cat# 017-000-121, West Grove, PA) and 0.3% Triton X-100 for 2 hours. Sections were then incubated for 24 hours at 4 °C in mouse anti-GAD67 (1:1000; Millipore Sigma, cat# MAB5406), rabbit anti-NeuN (1:500; Millipore Sigma, cat# ABN78), rabbit Neuro-Chrom pan neuronal marker (1:1000; Millipore-Sigma, cat# ABN2300), rabbit anti-NPY (1:1000; Peninsula Labs, cat# T-4070), guinea pig anti-GlyT2 (1:2000; Synaptic Systems cat# 272-004) or mouse anti-bNOS (1:1000, Millipore Sigma, cat#: N2280). On the following day, sections were rinsed in 0.1 M PBS and incubated in Alexa Fluor 647-tagged goat anti-mouse IgG or Alexa Fluor 750-tagged goat anti-rabbit IgG (1:100, Thermo Fisher, cat# A21235 and A21039), or Alexa Fluor 647-tagged donkey anti-mouse IgG, Alexa Fluor 647-tagged donkey anti-rabbit IgG, or Alexa Fluor 594-tagged goat anti-guinea pig IgG (1:500, Thermo Fisher, cat# A-21202, A-21206 and A-11076) for 1.0 – 1.5 hours at room temperature. Sections were then mounted on Superfrost Plus microscope slides (Thermo Fisher, cat# 12-550-15) and coverslipped using Fluoromount-G (SouthernBiotech, cat# 0100–01) or DPX (Sigma-Aldrich cat# 06522). Images were collected using a 1.30 NA 40x oil-immersion objective or a 1.40 NA 63x oil-immersion objective on a Leica TCS SP8 laser scanning confocal microscope.

### Antibody validation

All antibodies used in this study, except anti-NPY, have been previously validated in the IC. To label GABAergic neurons, we used the mouse monoclonal anti-GAD67 antibody (Millipore Sigma, cat# MAB5406). This antibody was raised against the 67 kDa isoform of glutamic acid decarboxylase (GAD). The manufacturer reports that western blot analysis showed no cross-reactivity with the 65 kDa isoform of GAD (GAD65). This antibody has been used in several studies to identify GABAergic neurons in the IC (Ito et al., 2009; Mellott et al., 2014; Beebe et al., 2016; Goyer et al., 2019). To identify neurons, we performed anti-NeuN staining with a rabbit polyclonal antibody (Millipore Sigma, cat# ABN78). The manufacturer reports that anti-NeuN specifically recognizes the DNA-binding, neuron-specific protein NeuN. Previous studies used this antibody to identify neurons in the IC (Foster et al., 2014; Mellott et al., 2014; Beebe et al., 2016; Goyer et al., 2019). To define the borders of IC subdivisions, we performed a double staining using anti-GAD67 and anti-GlyT2 antibodies (Buentello et al., 2015). Guinea pig anti-GlyT2 antibody (Synaptic Systems, cat# 272 004) was used to identify glycine transporter 2, which is localized in axons. We also immunostained with anti-brain nitric oxide synthase (bNOS) to define the borders of the IC subdivisions (Millipore Sigma, cat# N2280). This antibody was raised against the IgG1 isotype from the NOS-B1 hybridoma. The manufacturer reports that anti-bNOS reacts specifically with nitric oxide synthase (NOS), derived from brain (bNOS, 150 – 160 kDa). Previous studies used anti-bNOS to delineate the borders of IC subdivisions in guinea pig and mouse (Coote and Rees, 2008; Keesom et al., 2018). Finally, to label NPY, we used the polyclonal rabbit anti-NPY antibody (Peninsula Laboratories, cat# T-4070). Pretreatment with antiserum containing NPY peptide abolished all staining, reinforcing the high specificity of this antibody (Rowan et al., 1993). This antibody has been successfully used in cortex (Rose et al., 2009; Milstein et al., 2015) and hippocampus (Ledoux et al., 2009).

### Analysis of GAD67 staining

Analysis of GAD67 immunostaining was performed as previously reported (Goyer et al., 2019). Images from representative sections of the IC (n = 2 male P74 mice, three sections per mouse, one caudal, one medial and one rostral) were collected at 2 µm depth intervals with a 1.30 NA 40x oil-immersion objective on a Leica TCS SP8 laser scanning confocal microscope. Images were analyzed using Leica LAS X software (version 3.3.0). As previously reported, anti-GAD67 antibody did not penetrate the entire depth of the tissue sections (Beebe et al., 2016; Goyer et al., 2019). For this reason, the analysis was restricted to the top 10 – 12 µm of each section, where the antibody fully penetrated. Within this region, we manually marked every hrGFP^+^ cell body in the left IC. We analyzed the green (hrGFP) and red (Alexa 647) channels separately, so that labeling in one channel did not influence analysis of the other channel. After the hrGFP cells were marked, the red and green color channels were merged, and in every instance where a cell body contained markers for hrGFP, we evaluated if GAD67 was also present. The number of double-labeled cells was compared to the total number of hrGFP^+^ neurons to determine the percentage of hrGFP^+^ neurons that were GAD67^+^ as well.

### Analysis of NeuN staining with design-based stereology

We used a design-based stereology approach to estimate the numbers of hrGFP^+^ and NeuN^+^ neurons in brain sections that were stained with anti-NeuN antibody. This approach has been previously described and validated in the IC (Goyer et al., 2019). In brief, we collected systematic random samples using a virtual grid of 370 µm x 370 µm squares that was overlaid on the IC section. The starting coordinates for the grid were set using the Mersenne Twister random number generator in Igor Pro 7 or 8 (WaveMetrics Inc). We then collected images of the coordinates determined by the upper-left intersection of each grid-square that fell over the IC. Each image consisted of a 184 µm x 184 µm Z-stack collected at 1 µm depth intervals with a 1.40 NA 63x oil immersion objective on a Leica TCS SP8 confocal microscope. Eight to twenty images were collected per slice. Three slices (caudal, middle and rostral) were analyzed per mouse (n = 2). Using Neurolucida 360 (MBF Bioscience), we counted neurons using the optical fractionator approach (West et al., 1991). In this approach, we defined guard zones as ≥2 µm-thick regions at the top and bottom of the slice and excluded these from subsequent analysis. Neurons between the guard zones, within a 15 µm-thick region at the center of the slice, were counted by making a single mark at the top of each cell. Cells crossing the right and top borders of the image stack were counted, whereas those crossing the left and bottom borders were not. The red (NeuN) and green (hrGFP) color channels were analyzed separately, so that labeling in one channel did not influence analysis of the other. In all neurons counted, hrGFP^+^ cells were also NeuN^+^ (1097/1097 cells). The total number of double-labeled (hrGFP^+^/NeuN^+^) cells was then compared to the total number of NeuN^+^ cells. We also collected tile scan images of each IC section analyzed using a 10x objective and used GAD67 and GlyT2 staining to determine the border separating the ICc from the IC shell (ICs) regions. The 63x Z-stacks were aligned to these tile scans using Adobe Photoshop, and counted neurons were assigned to the ICc or IC shell.

### Analysis of GAD67 staining in Npy1r^cre^ mice with design-based stereology

To evaluate the neurotransmitter content of neurons that express the Y_1_R, we performed anti-GAD67 staining on brain slices of Npy1r^cre^ x Ai14 mice. We then used design-based stereology to quantify the proportion of tdTomato^+^ neurons that co-labeled with GAD67. This analysis was performed as described above, except that since anti-GAD67 antibody does not penetrate the entire depth of tissue sections (Beebe et al., 2016; Goyer et al., 2019), the analysis was restricted to depths at which clear anti-GAD67 immunostaining was present.

### Brain slice preparation

To characterize the intrinsic physiology of NPY neurons and the effect of NPY on neurons in the IC, whole-cell patch-clamp recordings were performed in NPY-hrGFP (van den Pol et al., 2009) and Npy1r^cre^ mice (Padilla et al., 2016). Both males (n = 22 for NPY-hrGFP mice, n = 11 for Npy1r^cre^ x Ai14 mice) and females (n = 33 for NPY-hrGFP mice, n = 10 for Npy1r^cre^ x Ai14 mice) aged P22 – P50 were used. No differences were observed between animals from different sexes (linear regression, p>0.05). Mice were deeply anesthetized with isoflurane and then rapidly decapitated. The brain was removed, and the IC was dissected in ∼34 °C artificial cerebrospinal fluid (ACSF) containing (in mM): 125 NaCl, 12.5 glucose, 25 NaHCO_3_, 3 KCl, 1.25 NaH_2_PO_4_, 1.5 CaCl_2_, 1 MgSO_4_, 3 sodium pyruvate and 0.40 L-ascorbic acid (Acros Organics), bubbled to a pH of 7.4 with 5% CO_2_ in 95% O_2_. Coronal sections of the IC (200 µm) were cut in ∼34 °C ACSF with a vibrating microtome (VT1200S, Leica Biosystems) and incubated at 34 °C for 30 min in ACSF bubbled with 5% CO_2_ in 95% O_2_. After incubation, slices were placed at room temperature for at least 30 min before being transferred to the recording chamber. All chemicals were obtained from Thermo Fisher Scientific unless otherwise noted.

### Electrophysiological recordings

For recordings, slices were placed into a chamber that was continuously perfused at ∼2 ml/min with 34°C oxygenated ACSF. NPY-hrGFP neurons and Y_1_R-tdTomato neurons in IC slices were identified with epifluorescence using a Nikon FN1 microscope. Whole-cell current-clamp recordings were performed with a BVC-700A patch clamp amplifier (Dagan Corporation). Data were low-pass filtered at 10 kHz, sampled at 50 kHz with a National Instruments PCIe-6343 data acquisition board, and acquired using custom software written in IgorPro.

Recording pipettes were pulled from borosilicate glass pipettes (outer diameter 1.5 mm, inner diameter 0.86 mm, cat# BF150-86-10, Sutter Instrument) using a P-1000 microelectrode puller (Sutter Instrument). Pipettes were filled with an internal solution containing (in mM): 115 K-gluconate, 7.73 KCl, 0.5 EGTA, 10 HEPES, 10 Na_2_phosphocreatine, 4 MgATP, 0.3 NaGTP, supplemented with 0.1% biocytin (w/v), pH adjusted to 7.3 with KOH and osmolality to 290 mmol/kg with sucrose. Pipettes with resistances of 3.0 - 4.8 MΩ when filled with internal solution were used for recordings.

Input resistance was assessed by applying a series of 100 ms current steps that hyperpolarized the membrane potential from just below rest to < −100 mV. For each step response, the peak and steady-state voltage changes were measured, and these values were used to generate voltage versus current plots from which the peak (*R*_*pk*_) and steady-state (*R*_*ss*_) input resistances were calculated based on the slope of a linear regression. Membrane time constant was determined by applying fifty, 300 ms current steps that hyperpolarized the membrane potential by 2 - 4 mV and taking the median of the time constants obtained by fitting an exponential function to each response. All membrane potential values were corrected for the liquid junction potential (−11 mV). Capacitance and series resistance were compensated using bridge balance. Recordings with series resistance >20 MΩ or changes in series resistance >15% during the recording period were discarded.

To test if NPY neurons fire in the presence of synaptic blockers (n = 8), we blocked synaptic inputs using 50 µM D-AP5 (NMDA receptor antagonist, Hello Bio, cat# HB0225) and 10 µM NBQX disodium salt (AMPA receptor antagonist, Hello Bio, cat# HB0443). All drugs were diluted in standard ACSF.

To evaluate the effects of Y_1_R activation, Y_1_R-tdTomato neurons were targeted for recordings. Since most NPY neurons spontaneously fire (n = 87 out of 121 cells), we included 1 µM tetrodotoxin (TTX, Hello Bio, cat# HB1035) in the ACSF in this experiment to block action potential mediated release of NPY. In current clamp mode, Y_1_R neurons were held for 10 min before starting the control period recording to limit possible effects of intracellular dialysis. During these 10 min, *R*_*pk*_, *R*_*ss*_ and membrane time constant were collected to monitor the health of the neuron. At the start of the 5 min control period, we initiated a protocol that recorded the resting membrane potential (*V*_*rest*_) every 10 seconds. Only neurons with a stable *V*_*rest*_, defined as *V*_*rest*_ drift <3 mV, were included in the analysis. We then applied 500 nM [Leu^31^, Pro^34^]-NPY (Tocris, cat # 1176), a high affinity Y_1_R agonist. We used 500 nM [Leu^31^, Pro^34^]-NPY based on previous studies performed in the hypothalamus (Roa and Herbison, 2012) and hippocampus (Giesbrecht et al., 2010). [Leu^31^, Pro^34^]-NPY was diluted in 10 ml of ACSF and bath applied for ∼5 min. The recording was then continued for a 20 - 50 min washout period. Only neurons that had a washout period over 20 min were included in the analysis. Control experiments followed the same approach, using a vehicle solution (normal ACSF).

To analyze the effect of [Leu^31^, Pro^34^]-NPY on the *V*_*rest*_ of IC neurons, we compared *V*_*rest*_ during the final two min of the baseline period with the initial 2 min of the drug application period (see Figure 7B, C, gray bars). The drug application period started 90 s after the end of the control period to allow time for the drug solution to reach the recording chamber. A cell was considered sensitive to [Leu^31^, Pro^34^]-NPY if it exhibited a >2 mV change in *V*_*rest*_ compared to baseline.

To extend the results from the bath application of [Leu^31^, Pro^34^]-NPY, in a subsequent experiment, [Leu^31^, Pro^34^]-NPY was delivered using a puffer pipette. The puffer pipette was placed close to the recorded cell, allowing the drug to reach Y_1_R neurons faster than was possible with the bath application. The puffer pipette was filled with 1 µM [Leu^31^, Pro^34^]-NPY diluted in ACSF. In current clamp mode, R_pk_, R_ss_ and membrane time constant were recorded for each cell. Next, we recorded 30 seconds of baseline and then puff applied [Leu^31^, Pro^34^]-NPY for 60 seconds (Acuna-Goycolea and van den Pol, 2005) (n = 6 animals, 9 cells). For control experiments (n = 5 animals, 7 cells), the puffer pipette was filled with ACSF and the position of the pipette and duration of the puff were identical to the drug group. To analyze the effect of puffing [Leu^31^, Pro^34^]-NPY, we compared the mean *V*_*rest*_ during the final 20 seconds of the baseline period with the mean *V*_*rest*_ during the middle 20 seconds of the puff application (20 – 40 seconds after puff onset).

### Post hoc reconstructions of morphology and morphology analysis

Neurons were filled with biocytin via the recording pipette. After the recording, an outside-out patch was formed, by slowly removing the pipette, to allow the cell membrane to reseal. Brain slices were then fixed overnight in 4% paraformaldehyde (PFA) in 0.1 M phosphate buffer (PB, pH 7.4) and moved to 0.1 M PBS the next day. Slices were stored at 4 °C for up to three weeks and then stained using fluorescent biocytin-streptavidin histochemistry. For this, slices were washed in 0.1 M PBS three times for 10 min (3 × 10 min in PBS), the membrane was permeabilized in 0.2% Triton X-100 in 0.1 M PBS for 2 hours, washed 3 × 10 min in PBS, and stained at 4 °C for 24 hours with streptavidin-Alexa Fluor 647 (1:1000). The next day, slices were washed 3 × 10 min with PBS, post fixed in 4% paraformaldehyde (PFA) in 0.1 M phosphate buffer (PB, pH 7.4) for 1 hour and then washed 3 × 10 min with PBS. Slices were then mounted on Superfrost Plus microscope slides in anti-fade media (Fluoromount-G). Images were obtained with a Leica TCS SP8 laser scanning confocal microscope using a 1.40 NA 63x oil-immersion objective. Z-stack tile scans were performed to image the soma and dendritic arbor. Image stacks were imported into Neurolucida 360 (MBF Bioscience), where three-dimensional reconstructions and quantitative analyses of neuronal morphology were performed.

### Retrograde tracing

In seven NPY-hrGFP mice, we used red fluorescent RetroBeads to evaluate whether NPY neurons participate in the IC-MG projection (“red beads,” Luma-Fluor, Inc., Naples, FL, USA; 1:2 dilution). Surgeries were performed on males and females, aged P22 – P28, using standard aseptic techniques. Mice were deeply anesthetized with isoflurane (2 – 3%) and their body temperature maintained with a homeothermic heating pad. The analgesic carprofen (5 mg/ kg, CarproJect, Henry Schein Animal Health) was injected subcutaneously. The scalp was shaved and a rostro-caudal incision was made along the midline to expose the skull. A unilateral craniotomy was performed, using a micromotor drill (K.1050, Foredom Electric Co.) with a 0.5 mm burr (Fine Science Tools). The coordinates were defined relative to the bregma suture, with injection depths relative to the surface of the brain. Injections were made in one penetration: 3200 µm caudal, 1900 µm lateral, and 2750 µm deep.

RetroBeads were injected with a NanoJect III nanoliter injector (Drummond Scientific Company) connected to a MP-285 micromanipulator (Sutter Instruments). Glass injection pipettes were pulled using 1.14 mm outer diameter, 0.53 mm inner diameter capillary glass (cat# 3-000-203-G/X, Drummond Scientific Company) with a P-1000 microelectrode puller (Sutter Instrument). The injector tip was cut and back-filled with mineral oil and then front-filled with the RetroBeads. After the injections were complete, the scalp was sutured, using Ethilon 6–0 (0.7 metric) nylon sutures (Ethicon USA LLC), and the incision was treated with 0.5 ml 2% Lidocaine hydrochloride jelly (Akorn Inc.). After recovery from anesthesia, mice were returned to the vivarium and were monitored daily until the wound healed. After 5 – 7 days, mice were perfused and projections were analyzed.

### Subdivisions of the IC

We used two different approaches to define the borders between the ICc, ICd, and IClc. For the stereology experiments and for the analysis of the distribution of NPY neurons, subdivision borders were defined using a combination of GlyT2 and GAD67 staining (Buentello et al., 2015). To identify the subdivisions in which reconstructed neurons were located, we used bNOS staining (Coote and Rees, 2008). Both methods have been previously validated in the IC, and we did not observe any obvious difference between the subdivision borders obtained with these approaches.

### Statistics

All statistical analyses were performed in Igor Pro. Data are presented as mean ± SD. The statistical tests used for each experiment are indicated in the results. Results were considered significant when P < 0.05. For comparing the intrinsic physiology of sustained and adapting Y_1_R-expressing neurons, the significance level (α) was adjusted with the Bonferroni correction to account for the multiple comparisons performed (adjusted α = 0.05/8 = 0.00625; Figure 7G).

## Results

### NPY neurons are distributed throughout the main subdivisions of the IC

To target experiments to NPY-expressing neurons, we obtained the NPY-hrGFP mouse line, in which hrGFP expression is driven by the NPY promoter (van den Pol et al., 2009). We performed immunostaining against the NPY peptide to determine whether this mouse line selectively labels NPY-expressing neurons in the IC. In brain slices from two P58 males, we counted hrGFP^+^ neurons in one caudal, one middle and one rostral section of the IC per animal (n = 3205 hrGFP^+^ neurons, Figure 1A-C). We then evaluated whether each hrGFP^+^ neuron co-labeled with anti-NPY. We found that 94.7% of hrGFP^+^ neurons were NPY^+^, indicating that the NPY-hrGFP mouse line provides highly selective labeling of NPY-expressing neurons (n = 3035 hrGFP^+^/NPY^+^ neurons and 170 hrGFP^+^/NPY^-^ neurons, Table 1).

**Figure 1.**
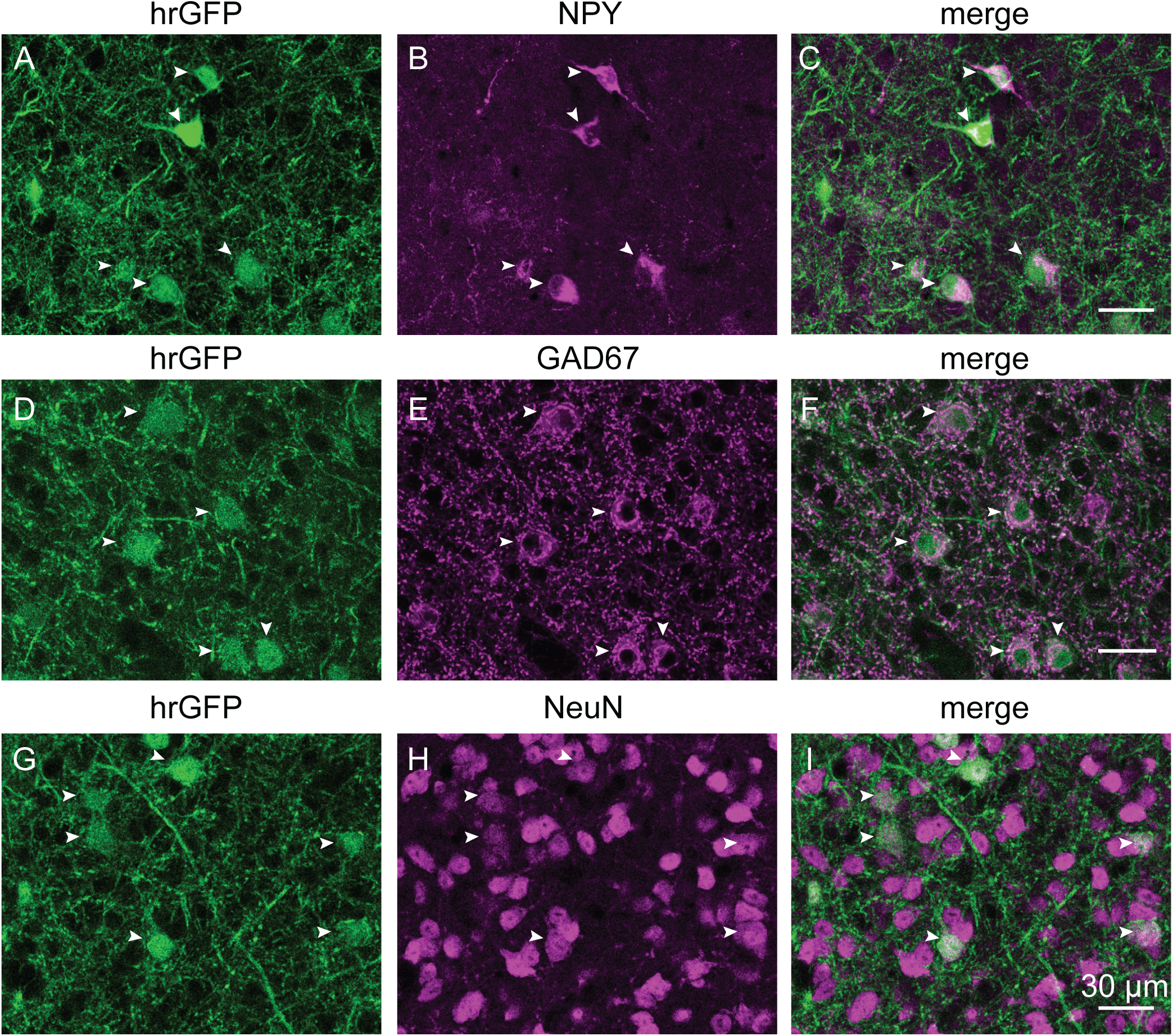
NPY neurons express NPY, are GABAergic, and represent 7.6% of neurons in the ICc. ***A, D, G***, Confocal images of NPY neurons expressing hrGFP in coronal sections of the ICc from NPY-hrGFP mice. ***B***, Confocal images showing immunostaining for NPY (magenta). ***C***, Nearly all hrGFP^+^ neurons colocalized with anti-NPY. ***E***, Confocal images showing immunostaining for GAD67 (magenta). ***F***, Nearly all hrGFP^+^ neurons were positive for GAD67 (magenta). ***H***, Confocal images showing immunostaining for NeuN (magenta). ***I***, All NPY neurons were positive for NeuN. White arrowheads indicate NPY neurons labeled by NPY, GAD67 and NeuN. Scale bar = 30 µm, applies to all images.

**Table 1.** hrGFP^+^ neurons express NPY. Across two mice, an average of 94.7% of hrGFP^+^ neurons were labeled with an antibody against NPY.

| Animal | Slice # | # hrGFP<sup>+</sup> | # NPY<sup>+</sup> | # hrGFP<sup>+</sup> NPY<sup>-</sup> | % hrGFP<sup>+</sup> NPY<sup>+</sup> |
| --- | --- | --- | --- | --- | --- |
| P58 male #1 | 1 (caudal) | 464 | 436 | 28 | 94.0 |
|  | 2 (middle) | 563 | 537 | 26 | 95.4 |
|  | 3 (rostral) | 567 | 537 | 30 | 94.5 |
|  | **Total** | **1594** | **1510** | **84** | **94.7** |
| P58 male #2 | 1 (caudal) | 496 | 485 | 11 | 97.8 |
|  | 2 (middle) | 602 | 555 | 47 | 92.2 |
|  | 3 (rostral) | 514 | 486 | 28 | 94.6 |
|  | **Total** | **1612** | **1526** | **86** | **94.7** |
| **Grand total** |  | **3206** | **3036** | **170** | **94.7** |
| **Average across two mice** |  |  |  |  | **94.7** |

We observed hrGFP-labeled cells throughout the rostro-caudal extent of the auditory midbrain in hrGFP-NPY mice, including a presence across IC subdivisions and in the nucleus of the brachium of the IC and the intercollicular tegmentum. We observed virtually no NPY neurons in auditory brainstem nuclei outside the midbrain. Within the auditory midbrain, about 95% of the NPY neurons were present in the three main subdivisions of the IC. Figure 2A shows the distribution of NPY neurons (green) through the rostro-caudal extent of the three major IC subdivisions. We plotted each individual NPY neuron in the IC in three cases (Figure 2B). The ICd contained the most NPY cells (46% ± 6% of the IC NPY population), while the IClc contained the fewest (13% ± 3%; Figure 2C). The ICd also had the highest average density of NPY neurons (288 ± 32 cells per mm^2^), while the IClc had the lowest average density (177 ± 28 cells per mm^2^; Figure 2D).

**Figure 2.**
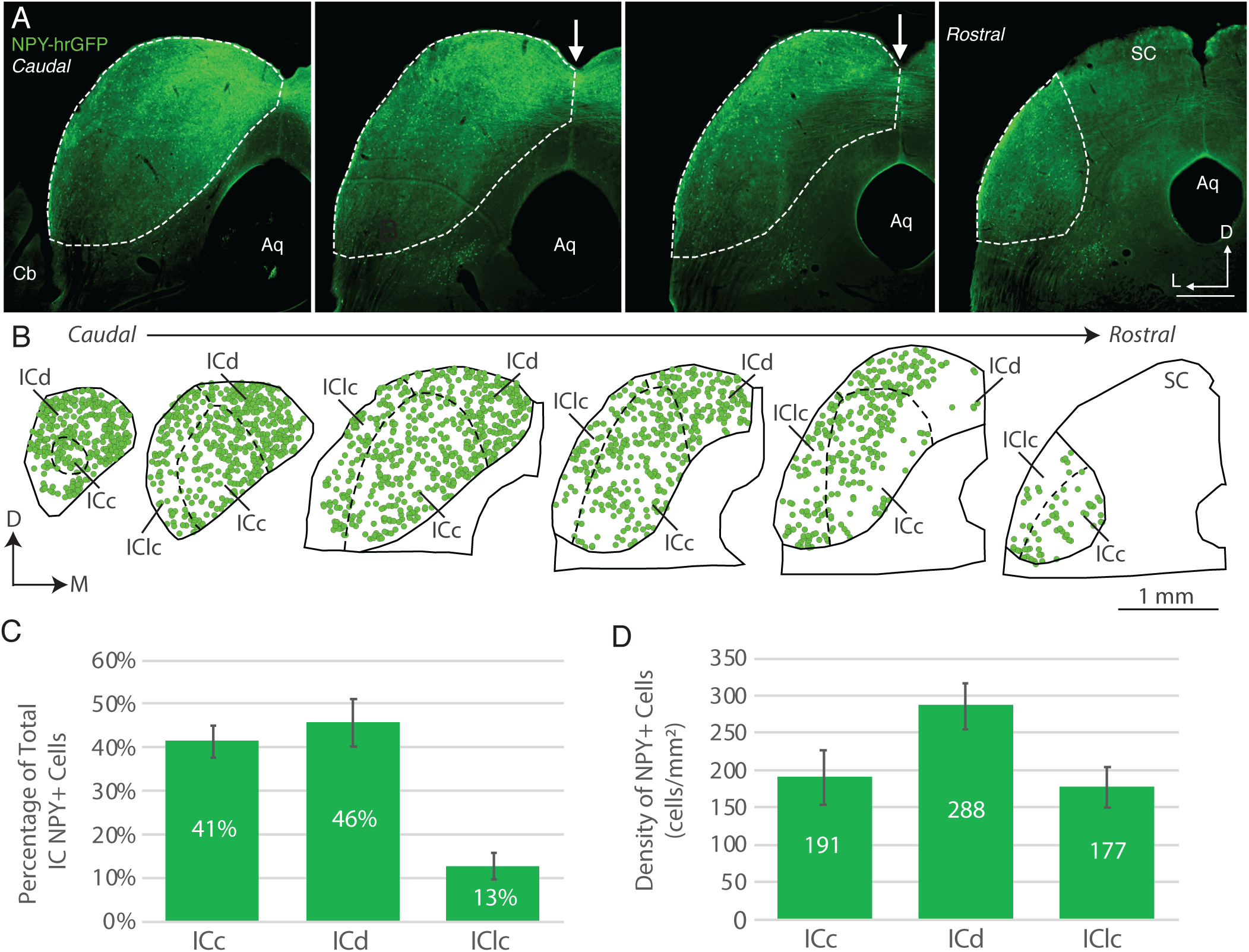
NPY neurons are distributed throughout the IC. ***A***, Micrographs of a series of coronal sections through the IC of an NPY-hrGFP mouse. NPY^+^ cells (green) were observed in all sections through the IC. White arrows indicate NPY^+^ axons crossing the IC commissure. Up is dorsal and left is lateral. Aq- cerebral aqueduct, Cb- cerebellum, SC- superior colliculus. Scale bar = 500 µm. ***B***, A plot through the three main subdivisions of the IC showing the distribution of NPY cells (each green circle represents one NPY neuron). Sections are arranged from caudal to rostral. Up is dorsal and right is medial. Scale bar = 1 mm. ***C***, A bar graph showing the mean ± SD of the proportion of the NPY^+^ population present in each of the main IC subdivisions. The ICd contained the highest proportion of the NPY^+^ population while the IClc contained the lowest proportion. ***D***, A bar graph showing the mean ± SD of the density of NPY^+^ cells in each of the IC subdivisions. The ICd contained the highest density of NPY^+^ cells and the IClc contained the lowest density.

### NPY neurons represent one-third of the GABAergic neurons in the IC

NPY neurons are nearly exclusively GABAergic in the cerebral cortex (Karagiannis et al., 2009), hippocampus (Milner et al., 1997) and hypothalamus (Henry et al., 2015; Marshall et al., 2017). To investigate the neurotransmitter content of NPY neurons in the IC, we performed immunostaining against GAD67, an enzyme essential for GABA synthesis. Using brain slices from two P74 males (Figure 1D-F), we counted hrGFP^+^ neurons in one caudal, one middle and one rostral section of the IC per animal (n = 2673 hrGFP^+^ neurons). We then determined whether each hrGFP^+^ neuron co-labeled with anti-GAD67. Out of the 2673 hrGFP^+^ neurons counted, 98.5% were GAD67^+^ (n = 2633 hrGFP^+^/ GAD67^+^ neurons and 40 hrGFP^+^/GAD67^-^ neurons, Table 2). We suspect that the absence of GAD67 immunostaining in 40 hrGFP^+^ neurons was due to rare cases of poor antibody penetration or nonspecific expression of hrGFP. Thus, these data indicate that NPY neurons in the IC are GABAergic.

**Table 2.** NPY neurons are GABAergic. Across two mice, an average of 98.5% of hrGFP^+^ neurons were labeled with an antibody against GAD67.

| Animal | Slice # | # hrGFP<sup>+</sup> | # GAD67<sup>+</sup> | # hrGFP<sup>+</sup> GAD67<sup>-</sup> | % hrGFP<sup>+</sup> GAD67<sup>+</sup> |
| --- | --- | --- | --- | --- | --- |
| P74 male #1 | 1 (caudal) | 456 | 450 | 6 | 98.7 |
|  | 2 (middle) | 387 | 380 | 7 | 98.2 |
|  | 3 (rostral) | 328 | 322 | 6 | 98.2 |
|  | **Total** | **1171** | **1152** | **19** | **98.4** |
| P74 male #2 | 1 (caudal) | 468 | 462 | 6 | 98.7 |
|  | 2 (middle) | 346 | 337 | 9 | 97.4 |
|  | 3 (rostral) | 688 | 682 | 6 | 99.1 |
|  | **Total** | **1502** | **1481** | **21** | **98.6** |
| **Grand total** |  | **2673** | **2633** | **40** | **98.5** |
| **Average across two mice** |  |  |  |  | **98.5** |

Next, we used design-based stereology to determine the percentage of neurons in the IC that express NPY. For this experiment, we analyzed the ICc and IC shell (ICs = ICd plus IClc) separately, defining the borders between these subdivisions according to the pattern of GAD67 and GlyT2 immunoreactivity (Buentello et al., 2015). Sections from two P54 male NPY-hrGFP mice were stained with anti-NeuN, a neuron-selective antibody previously shown to label most or all neurons in the IC (Figure 1G-I; Mellott et al., 2014; Beebe et al., 2016). Three sections per mouse were analyzed: one caudal, one middle, and one rostral. To ensure unbiased counting of neurons, we applied the optical fractionator method (see Materials and Methods). Sections adjacent to the anti-NeuN sections were immunostained with anti-GAD67 and anti-GlyT2, and IC subdivision boundaries were determined by comparing these to the adjacent anti-NeuN stained sections. This analysis showed that NPY neurons represented 7.6% of neurons in the ICc and 7.8% in the ICs (Table 3). Since previous studies indicate that 20 – 27% of IC neurons are GABAergic (Oliver et al., 1994; Merchán et al., 2005; Beebe et al., 2016), our results suggest that NPY-expressing neurons account for approximately one-third (27 – 39%) of GABAergic neurons in the IC.

**Table 3.** NPY neurons represent 7.6% of neurons in the ICc and 7.8% of neurons in the IC shell. Table shows stereological analysis of the percentage of neurons (NeuN^+^) in the ICc and IC shell (ICd + IClc) that expressed hrGFP in NPY-hrGFP mice.

| ICc |  |  |  |  |
| --- | --- | --- | --- | --- |
| Coronal Slice | P58 Male #1 | P58 Male #2 | Per slice pane | Grand average |
| Caudal | 8.0% | 9.4% | 8.2% |  |
|  | (60/755) | (15/159) | (75/914) |  |
|  | 7 samples | 4 samples | 11 samples |  |
| Medial | 7.9% | 7.0% | 7.4% |  |
|  | (84/1070) | (90/1278) | (174/2348) |  |
|  | 11 samples | 9 samples | 20 samples |  |
| Rostral | 7.7% | 7.2% | 7.5% |  |
|  | (77/999) | (68/942) | (145/1941) |  |
|  | 10 samples | 10 samples | 20 samples |  |
| Per mouse | 7.8% | 7.3% |  | 7.6% |
|  | (221/2824) | (173/2379) |  | (394/5203) |

| IC shell |  |  |  |  |
| --- | --- | --- | --- | --- |
| Coronal Slice | P58 Male #1 | P58 Male #2 | Per slice pane | Grand average |
| Caudal | 8.6% | 6.3% | 8.1% |  |
|  | (140/1620) | (29/460) | (169/2080) |  |
|  | 12 samples | 7 samples | 19 samples |  |
| Medial | 6.9% | 10.4% | 7.9% |  |
|  | (128/1849) | (76/728) | (204/2577) |  |
|  | 11 samples | 8 samples | 19 samples |  |
| Rostral | 8.3% | 6.7% | 7.2% |  |
|  | (55/668) | (77/1155) | (132/1823) |  |
|  | 12 samples | 11 samples | 23 samples |  |
| Per mouse | 7.8% | 7.8% |  | 7.8% |
|  | (323/4137) | (182/2343) |  | (505/6480) |

Together, the above data show that NPY is expressed in a population of GABAergic neurons in the IC, but do not indicate whether this population represents one or several neuron types. To test this, we next analyzed the physiological and morphological properties of NPY neurons.

### Most NPY neurons exhibit a sustained firing pattern and have a propensity to spontaneously fire

In acute brain slices, we targeted whole-cell current clamp recordings to NPY-hrGFP neurons in the ICc, ICd, and IClc. The location of the recording was noted at the end of the recording and confirmed during retrieval of neuronal morphology (morphology was recovered from 49 out of 146 recorded neurons). Because the intrinsic physiology of NPY neurons did not differ across IC subdivisions, neurons from ICc, ICd, and IClc are lumped together here. Most NPY neurons presented moderate voltage sag in response to hyperpolarizing current steps (Figure 3A, left) and a sustained firing pattern in response to depolarizing current steps (Figure 3A, right). Firing versus current functions were determined with 1 s depolarizing current steps and showed that NPY neurons had relatively linear input-output functions (Figure 3B). NPY neurons had an average resting membrane potential of -63.4 ± 3.8 mV (Figure 3C_1_) and exhibited relatively low expression of *I*_*h*_ current, with voltage sag ratios (steady-state/peak) of 0.89 ± 0.09 and 0.90 ± 0.09 measured from current steps that elicited peak hyperpolarization of 91.1 ± 1.8 mV and 111.0 ± 0.7 mV, respectively (Figure C_2_). In addition, NPY neurons exhibited moderate membrane time constants (15.9 ± 8.6 ms, Figure 3C_3_) and input resistances (*R*_*pk*_ = 238.7 ± 124.0 MΩ, *R*_*ss*_ = 249.5 ± 167.0 MΩ, Figure 3C_4_) and cells that did not spontaneously fire had low rheobase values (46.7 ± 25.1 pA, Figure 3C_5_).

**Figure 3.**
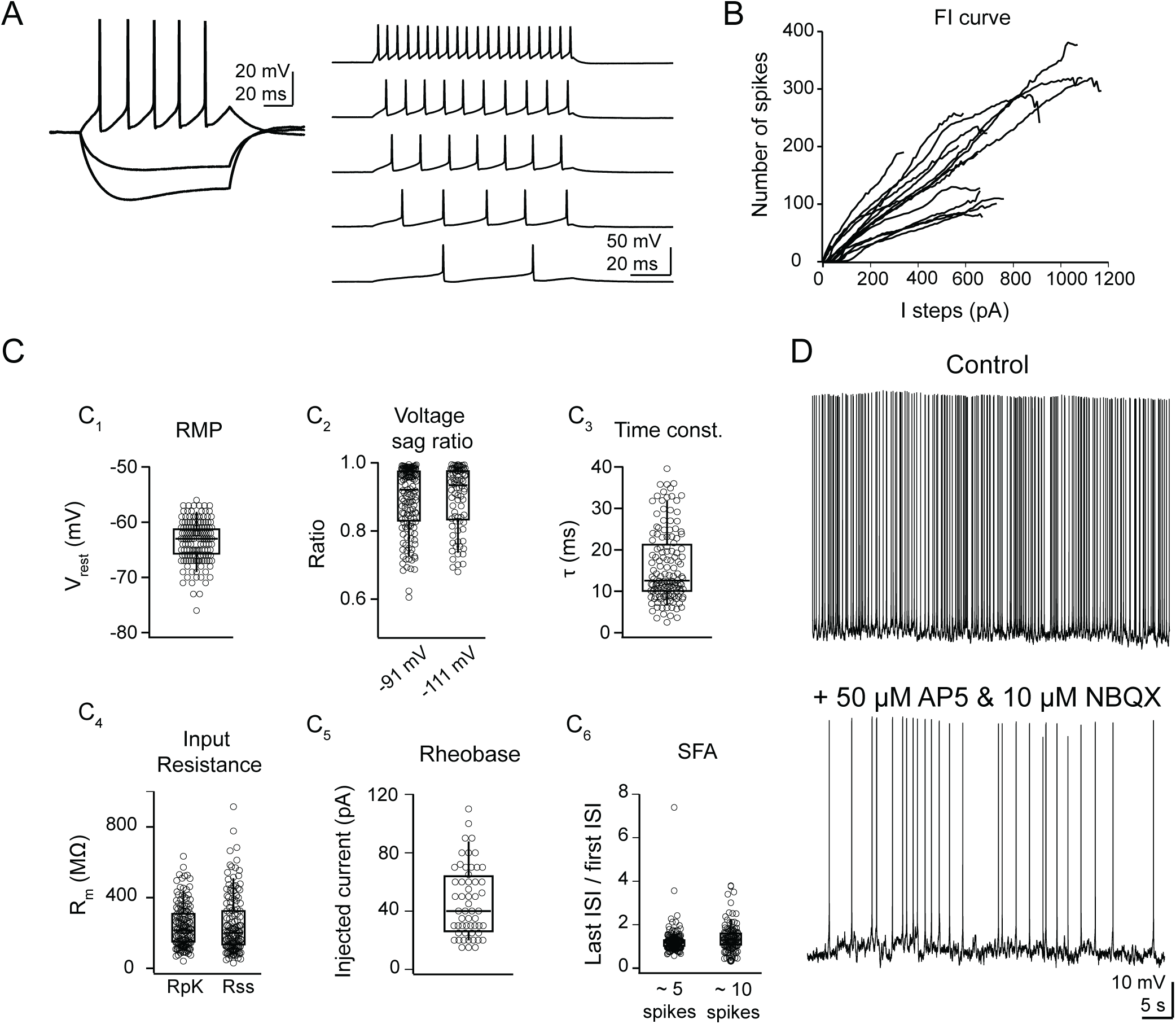
NPY neurons have a sustained firing pattern and most fire spontaneously. ***A***, NPY neurons exhibited a sustained firing pattern in response to depolarizing current steps and a moderate voltage sag in response to hyperpolarizing current steps. ***B***, Firing vs input analysis showed a linear input-output response, with increasing firing rates in response to increasingly large depolarizing current steps. Example FI curves from 15 neurons are shown. ***C***, NPY neurons exhibited moderate intrinsic physiological properties. ***C***_***1***_, resting membrane potential, ***C***_***2***_, voltage sag ratio, ***C***_***3***_, membrane time constant, ***C***_***4***_, input resistance, ***C***_***5***_, rheobase, and ***C***_***6***_, spike frequency adaptation ratio. Boxplots show median, 25^th^ and 75^th^ percentile (box), and 9^th^ and 91^st^ percentile (whiskers). ***D***, Most NPY neurons spontaneously fired at rest, and spontaneous firing persisted in 7 out of 8 cells in the presence of synaptic blockers (50 µM D-AP5 and 10 µM NBQX).

Neurons were classified as having a sustained firing pattern if their spike frequency adaptation ratio (SFA ratio, last interspike interval/first interspike interval) was less than 2 and as adapting if their SFA ratio was greater than 2 (Peruzzi et al., 2000). We found that 92.4% of NPY neurons (n = 135 out of 146) had a sustained firing pattern in response to current steps that elicited ∼5 spikes and 89.7% (n = 96 out of 107) had a sustained firing pattern in response to current steps that elicited ∼10 spikes (Figure 3C_6_). Because neurons in the IC exhibit diverse firing patterns, the finding that nearly all NPY neurons had sustained firing patterns supports the hypothesis that NPY neurons constitute a distinct neuron class.

Interestingly, 71.9% of NPY neurons fired spontaneously at rest, with a median firing rate of 0.22 Hz (n = 87 out of 121 neurons, range: 0.01 – 28.06 Hz, mean = 2.70 Hz; Figure 3D). To determine whether spontaneous firing was driven by spontaneous synaptic input, we applied 50 µM D-AP5 and 10 µM NBQX to block AMPA and NMDA receptors and found that spontaneous firing remained in 7 out of 8 neurons (Figure 3D). These results suggest that NPY neurons may able to provide a GABAergic and NPY tone in the IC.

### NPY neurons have stellate morphology

Using biocytin-streptavidin staining, we reconstructed the morphology of 49 NPY neurons. Twenty-four of these neurons were located in the ICc, 18 in ICd and 7 in IClc. Figure 4 shows examples of the morphology of NPY neurons in the ICc, ICd and IClc. For comparison purposes, neurons are oriented as they would appear in a coronal section of the left IC viewed from a caudal perspective. Visual inspection suggested that NPY neurons had a stellate morphology in all subdivisions. This is expected for ICd and IClc, where most neurons are stellate, but in the ICc, only 15 - 20% of neurons have stellate morphology (Oliver and Morest, 1984; Oliver et al., 1991).

**Figure 4.**
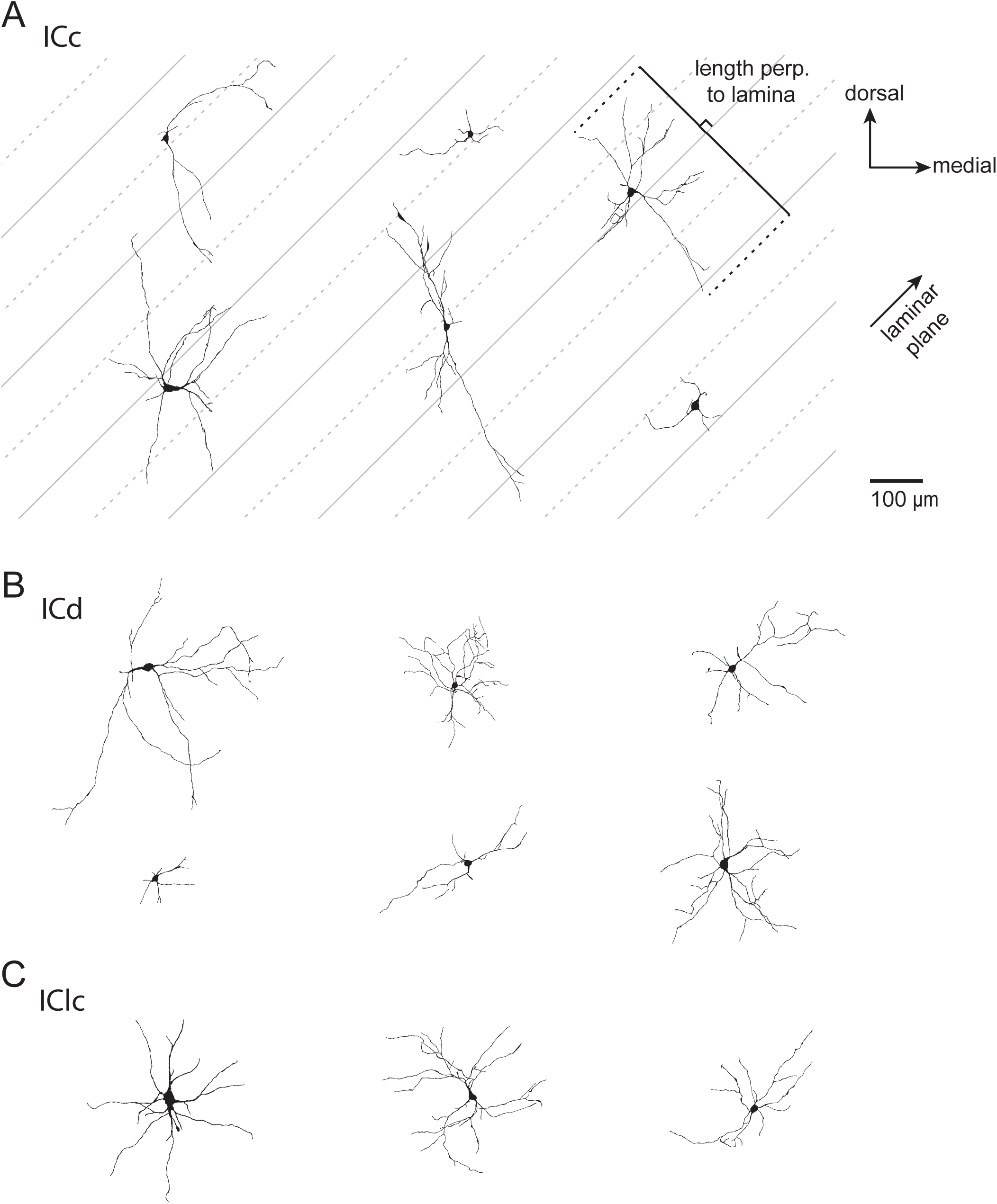
Morphology of NPY neurons in the ICc, ICd, and IClc. Representative reconstructions of the morphology of 6 NPY neurons from the ICc (***A***), 6 NPY neurons from the ICd (***B***), and 3 NPY neurons from the IClc (***C***). The gray lines in the ICc representation were drawn to illustrate the approximate orientation and spacing of the isofrequency laminae (45° angle). Solid gray lines are spaced 200 µm apart, dashed lines and solid lines are spaced 100 µm apart.

We used several quantitative approaches to test whether the dendritic arbors of NPY neurons in the ICc were consistent with the stellate morphology predicted by visual inspection. First, we used two-dimensional principal component analysis (PCA) to determine the orientation and extent of the “length” (first principal direction) and “width” (second principal direction) axes of each NPY neuron within the coronal plane (Figure 5A). In the mouse, the isofrequency laminae extend at a ∼40° - 50° angle relative to the horizontal plane, and it is expected that the long axis of disc-shaped neurons extends parallel to the isofrequency laminae (Stiebler and Ehret, 1985). Our data show that the long axis of NPY neurons in the ICc did not have a preferred orientation, and only 3 of 24 neurons had a long axis oriented between 30° and 60° (Figure 5B).

**Figure 5.**
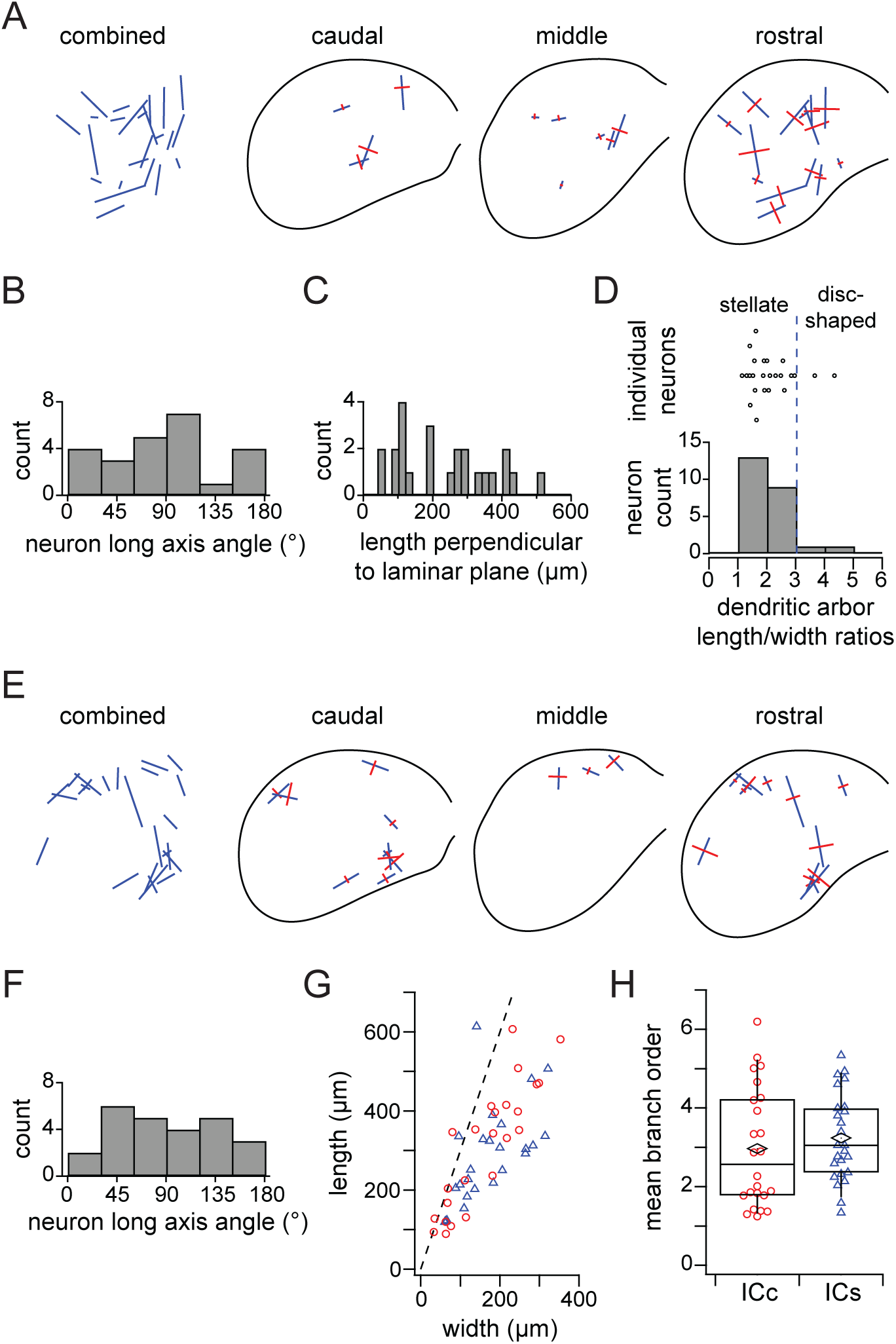
NPY neurons are a class of stellate cells. ***A***, Orientation of the dendritic arbors of NPY neurons from the ICc. Combined: all reconstructed neurons from the ICc (n = 24). Blue lines indicate the orientation and length of the longest axis (first determined using 2D PCA. Caudal, middle and rostral: orientation of dendritic fields separated according to position along the rostro-caudal axis of the ICc. Blue lines indicate longest axis, perpendicular red lines indicate second longest axis (second principal direction) as determined by 2D PCA. ***B***, Angular orientation of the long axis for every reconstructed NPY neuron located in the ICc. Angles indicate counter-clockwise rotation relative to the medial-lateral (horizontal) axis. ***C***, Extent of dendritic arbors of ICc NPY neurons perpendicular to the plane of the isofrequency laminae (45°). ***D***, 3D PCA was used to determine NPY neurons had a length to width ratio <3, indicating that they are stellate cells. ***E***, Orientation of the dendritic fields of NPY neurons from the ICd and IClc (combined as ICs). Colors of lines andaxis for every reconstructed NPY neuron located in the ICs (n = 25). Angles indicate counter-clockwise rotation relative to the medial-lateral (horizontal) axis. ***G***, Comparison of the mean dendritic branch order between NPY neurons in the ICc and ICs. ***H***, Comparison of the length/width ratio between NPY neurons in the ICc and ICs. The dashed line indicates length/width ratio = 3.

Second, because the dendrites of ICc stellate neurons typically extend across more than one isofrequency lamina, we measured how far ICc NPY neurons extended their dendrites perpendicular to a 45° laminar plane. Although the width of isofrequency laminae has not been determined in mice, the distance between laminae in rats presumably provides a conservative estimate for mice and ranges between 90 µm and 150 µm (Malmierca et al., 1993). We found that only 4 out of 24 ICc NPY neurons had dendritic arbors that extended <90 µm perpendicular to the laminar plane, and only 9 out of 24 had dendrites that extended <150 µm perpendicular to the laminar plane (Figure 5C). This suggests that the majority of ICc NPY neurons have dendritic arbors that extend across two or more isofrequency laminae.

Third, we performed three-dimensional principal component analysis on the x, y, z coordinate set for each ICc NPY neuron to determine the length (extent along the first principal direction) and width (extent along the second principal direction) of each neuron in three dimensions. We then calculated the length-to-width ratio of the dendritic arbor. Oliver and colleagues showed that stellate neurons have a length-to-width ratio <3 and disc-shaped neurons have a ratio ≥ 3 (Oliver et al., 1991). We found that 22 out of 24 NPY neurons (92%) in the ICc had a length to width ratio < 3, consistent with the hypothesis that NPY neurons are stellate (Figure 5D). Combined, the above quantitative analyses support the conclusion that ICc NPY neurons are a class of stellate cells. Because stellate cells represent 15 – 20% of ICc neurons (Oliver and Morest, 1984; Oliver et al., 1991) and our stereology data (Figure 2 G-I) indicate that NPY neurons represent 7.6% of neurons in the ICc, NPY neurons represent 38 - 50% of stellate neurons in the ICc.

We also performed two-dimensional principal component analysis on NPY neurons located in the ICd and IClc (combined as ICs; Figure 5E). As in the ICc, the longest axis of NPY neurons in the ICs did not exhibit a preferred orientation (Figure 5F). Instead, the long axes of ICs NPY neurons appeared to run parallel to the outer edge of the IC. Next, we compared the morphology of NPY neurons in the ICc versus those in the ICs. Interestingly, no difference was observed in the mean branch order number (ICc: 3.11 ± 1.54; ICs: 3.11 ± 1.04; two-tailed Mann-Whitney-Wilcoxon test, p = 0.713; Figure 5G) or the dendritic length/width ratio (ICc: 2.16 ± 0.87; ICs: 1.74 ± 0.55; two-tailed Mann-Whitney-Wilcoxon test, p = 0.091; Figure 5H). Thus, there were no gross morphological differences between NPY neurons in the ICc and ICs.

### NPY neurons project to the MG

Most IC neurons, including GABAergic IC neurons, are believed to project to targets outside the IC (Oliver et al., 1991). A major target of the GABAergic neurons is the ipsilateral MG (Peruzzi et al., 1997; Mellott et al., 2014). In order to investigate whether NPY neurons project within the IC-MG pathway, we placed red Retrobeads (RB) into the MG in NPY-hrGFP mice (Figure 6A). RB-labeled cells were numerous in the IC, and included cells that were double-labeled with RB and GFP, identifying NPY cells that project to the MG (Figure 6B,C, white arrows). Such double-labeled cells were present in all subdivisions of the ipsilateral IC. We also observed RB-labeled cells that were not NPY^+^ (Figure 6B, C, red arrows), indicating that there are many non-NPY^+^ cells within the IC-iMG pathway, consistent with previous reports that non-GABAergic (i.e., glutamatergic) IC cells also project to the MG. Finally, we observed NPY cells that did not contain RB (Figure 6B green arrows); these neurons could project to the MG but were not labeled by the MG injection, or they could represent IC neurons that project to a target other than the MG.

**Figure 6.**
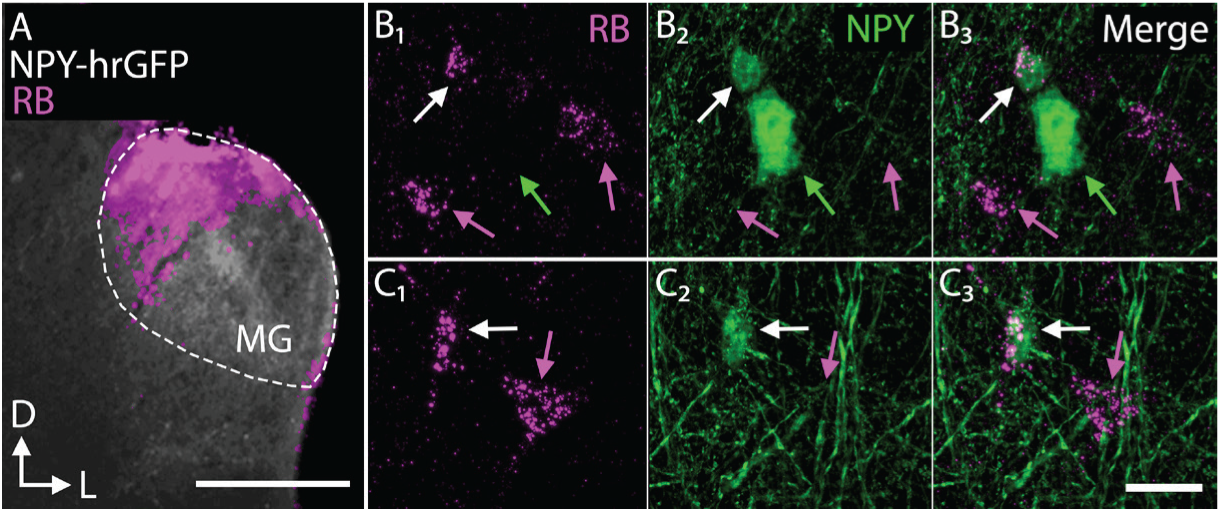
NPY neurons participate in the IC-MG pathway. ***A***, Low-magnification micrograph showing an injection of red Retrobeads (RB, red) into the MG of an hrGFP-NPY mouse. To aid in the visualization of the RB injection, hrGFP is shown in white. The approximate bounds of the MG are shown using a white dotted outline. Up is dorsal and right is lateral. Scale = 500 µm. ***B***,***C***, High-magnification photomicrographs showing double-labeled (RB-labeled/ NPY^+^) neurons (white arrows) in the IC ipsilateral to the injected MG. Nearby neurons could be single-labeled for RB (red arrows; B_1_, C_1_) or GFP (green arrows; B_2_, C_2_). Scale = 20 µm.

### Y_1_R neurons are glutamatergic neurons with heterogeneous intrinsic physiology

Previous studies suggest that Y_1_Rs are expressed in the IC (Kishi et al., 2005; Eva et al., 2006), but the neurotransmitter content and intrinsic physiology of the neurons expressing Y_1_Rs are unknown. We used Npy1r^cre^ x Ai14 mice to label Y_1_R-expressing neurons with the fluorescent protein tdTomato (Padilla et al., 2016) and found that Y_1_R neurons are broadly distributed in the IC (Figure 7 A-B). To investigate the neurotransmitter content of Y_1_R neurons, we performed immunofluorescence with anti-GAD67 on brain slices from two Npy1r^cre^ x Ai14 mice, aged P58. Using design-based stereology, we found that across 2148 tdTomato^+^ neurons, only 24 neurons co-labeled for GAD67 (1.1%, Table 4), suggesting that Y_1_R neurons in the IC are glutamatergic (Figure 7 C-E). Interestingly, we also observed that blood vessels in the IC were lined with non-neuronal cells expressing Y_1_R (Figure 7 A – E, blue arrowheads). Because NPY induces vasoconstriction in other brain regions (Bao et al., 1997; Cauli et al., 2004), NPY signaling might regulate blood flow in the IC, an important hypothesis we will pursue in a future study.

**Figure 7:**
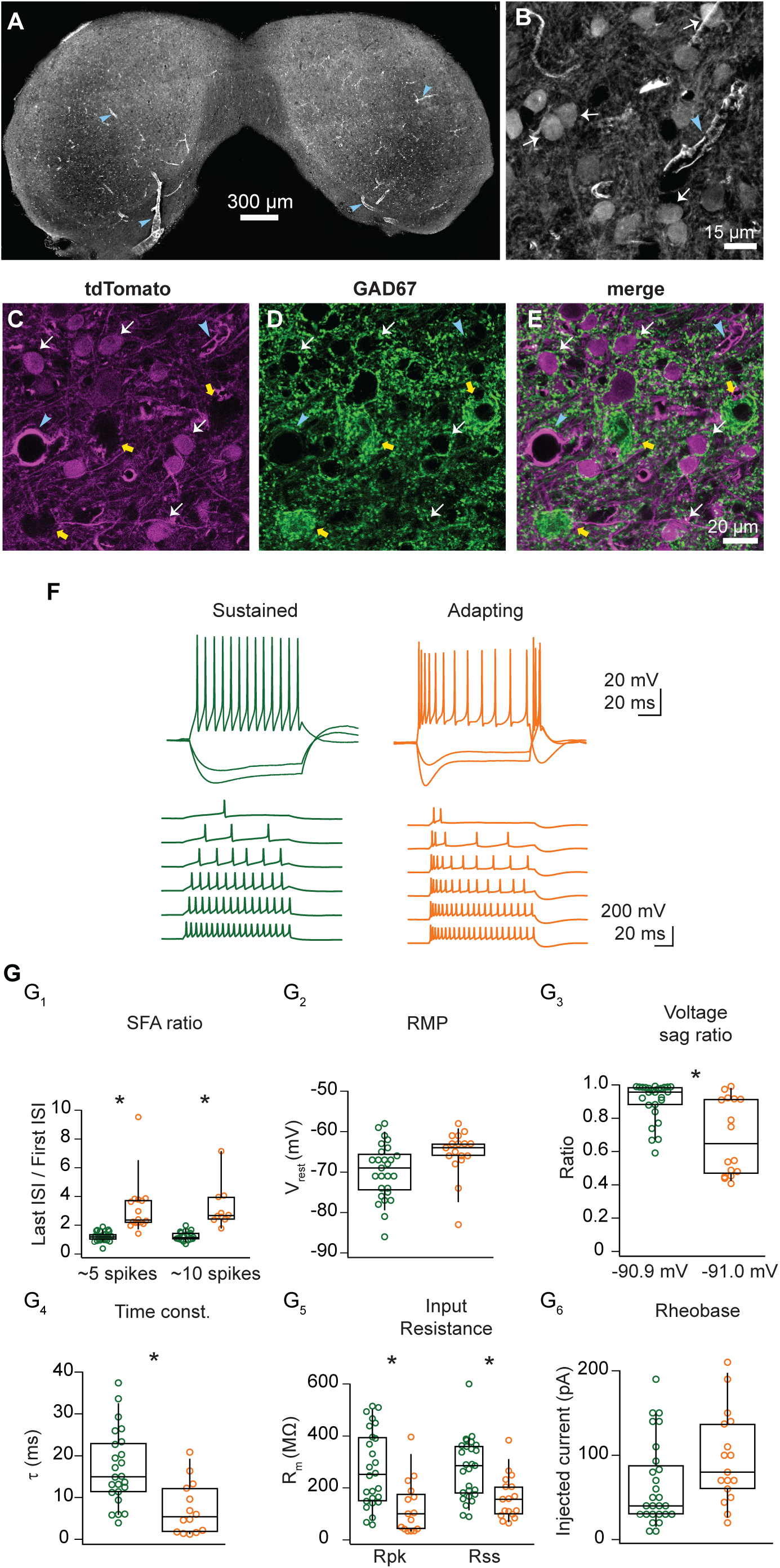
Y_1_R-expressing neurons are glutamatergic and exhibit sustained or adapting firing. ***A***, Tile scan image of tdTomato expression in a coronal section from an Npy1r^cre^ x Ai14 mouse. tdTomato^+^ neurons were broadly distributed throughout the IC. ***B***, Higher magnification image from the same section as ***A***. White arrows indicate cell bodies of neurons that were tdTomato^+^, indicating that they expressed Y_1_R. Blue arrows in ***A*** and ***B*** indicate blood vessels lined with tdTomato^+^ cells. ***C, D, E***, Confocal images of a coronal IC section from an Npy1r^cre^ x Ai14 mouse. tdTomato^+^ cells (***C***) did not co-label with GAD67 staining (***D***), suggesting the Y1R neurons are glutamatergic (***E***, merge). ***F***, Y_1_R neurons exhibited one of two different firing patterns in response to depolarizing current steps: sustained (left, green) or adapting (right, orange). ***G***, Sustained and adapting Y_1_R neurons exhibited heterogeneous intrinsic physiological properties. ***G***_***1***_, SFA ratio, ***G***_***2***_, resting membrane potential, ***G***_***3***_, voltage sag ratio, ***G***_***4***_, membrane time constant, ***G***_***5***_, input resistance, and ***G***_***6***_, rheobase. Boxplots show median, 25^th^ and 75^th^ percentile (box), and 9^th^ and 91^st^ percentile (whiskers). * = p < 0.00625.

**Figure 8.**
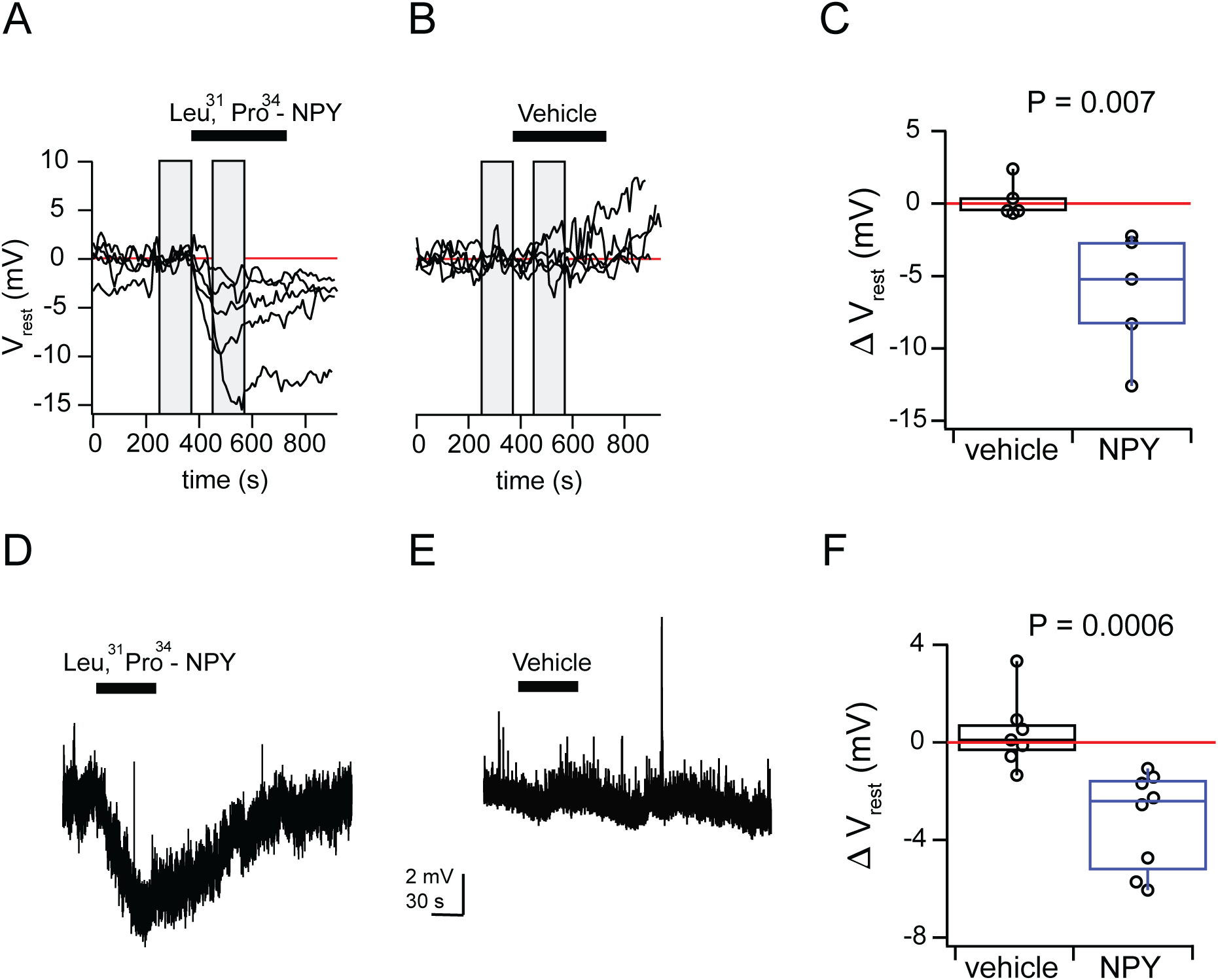
[Leu^31^, Pro^34^]-NPY hyperpolarized *V*_*rest*_ in a subset of Y_1_R-expressing IC neurons. ***A, B*,** Graphs showing 15 min of recording from tdTomato^+^ Y_1_R neurons. Gray bars indicate periods in which baseline (left bar) and treatment (right bar) measurements were made. ***A***, Bath application of 500 nM [Leu^31^, Pro^34^]-NPY led to a hyperpolarizing shift in the *V*_*rest*_ of 50% of tdTomato^+^ neurons (−6.2 ± 4.3 mV change; n = 5 out of 10 neurons; two-tailed paired t-test comparing baseline *V*_*rest*_ to treatment *V*_*rest*_, p = 0.032). ***B***, Vehicle had no effect on the membrane potential of tdTomato^+^ neurons (0.2 ± 1.3 mV change; n = 5; two-tailed paired t-test, p = 0.694). ***C***, Boxplot showing the change in *V*_*rest*_, between vehicle- and [Leu^31^, Pro^34^]-NPY-treated neurons (two-tailed Mann-Whitney-Wilcoxon test, p = 0.007). ***D***, Puff application of [Leu^31^, Pro^34^]-NPY led to a hyperpolarizing shift in the *V*_*rest*_ of 89% of tdTomato^+^ neurons (−3.1 ± 2.0 mV change; n = 8 out of 9 neurons; two-tailed paired t-test comparing baseline *V*_*rest*_ to treatment, p = 0.007). ***E***, Vehicle application had no effect on the membrane potential (0.4 ± 1.4 mV change; n = 7; two-tailed paired t-test comparing baseline *V*_*rest*_ to treatment, p = 0.57). ***F***, Boxplots showing the change in *V*_*rest*_ between vehicle and [Leu^31^, Pro^34^]-NPY-treated neurons (two-tailed Mann-Whitney-Wilcoxon test, p = 0.006). Boxplots show median, 25^th^ and 75^th^ percentile (box), and 9^th^ and 91^st^ percentile (whiskers).

**Table 4.**
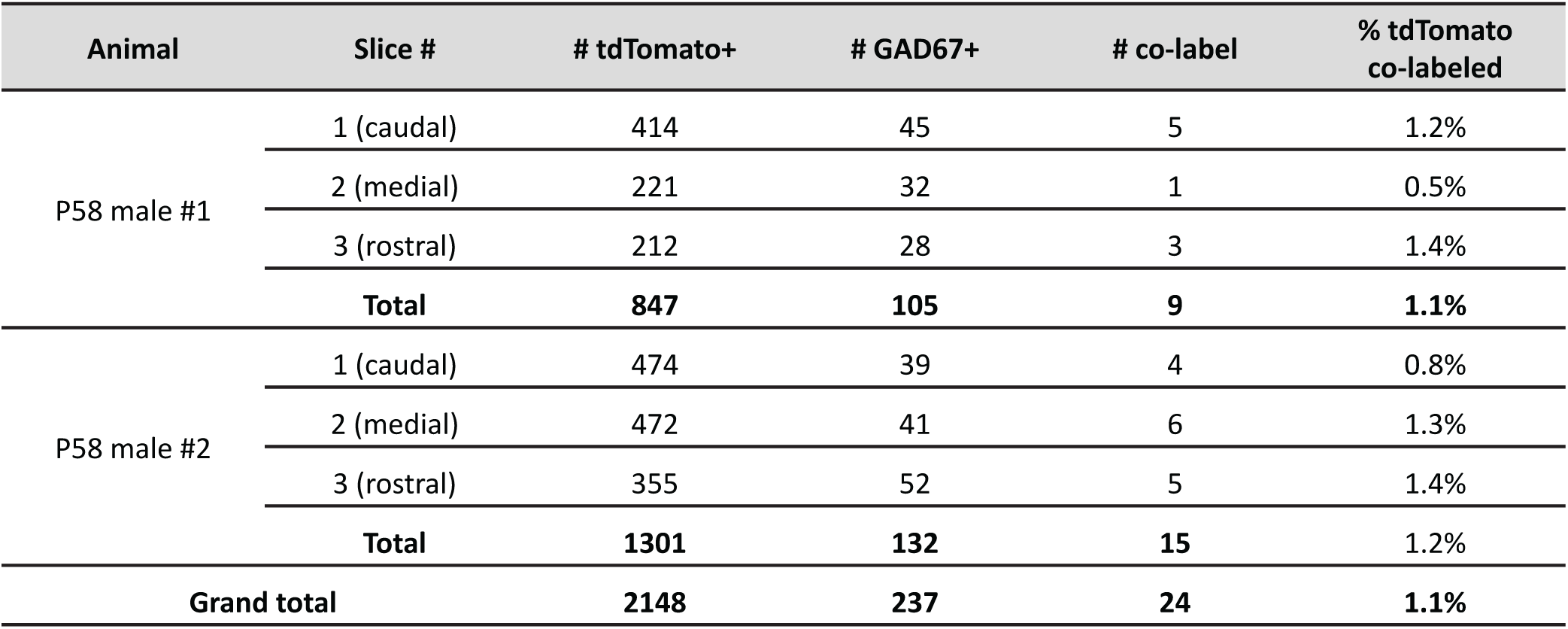
Y_1_R neurons are glutamatergic. Across two mice, an average of 1.1% of tdTomato^+^ neurons were labeled with an antibody against GAD67.

We next targeted whole-cell current clamp recordings to Y_1_R expressing neurons in acute brain slices from Npy1r^cre^ x Ai14 mice. In contrast to the rather homogenous intrinsic properties of NPY-hrGFP neurons, we found that Y_1_R neurons exhibited two distinct firing patterns that correlated with distinct intrinsic physiological properties. Slightly more than half of Y_1_R neurons exhibited a sustained firing pattern in response to depolarizing current steps, while the remaining Y_1_R neurons exhibited an adapting firing pattern (27 out of 44 neurons had a SFA ratio less than 2, and 17 out of 44 neurons had a SFA ratio greater than 2; Figure 7F and G_1;_ SFA ratio calculated with ∼5 spikes comparing sustained vs adapting neurons, two-tailed t-test, p = 0.0009; SFA ratio calculated with ∼10 spikes, two-tailed t-test, p = 0.002; Bonferroni corrected α = 0.00625). Sustained Y_1_R neurons had an average resting membrane potential of 69.8 ± 6.8 mV, while adapting Y_1_R neurons trended toward a more depolarized resting membrane potential of 65.3 ± 5.8 mV (Figure 7G_2_, two-tailed t-test, p = 0.026; Bonferroni corrected α = 0.00625). Adapting Y_1_R neurons had a more prominent voltage sag in response to hyperpolarizing current steps when compared to sustained neurons (Figure 7F, right), with voltage sag ratios (steady-state/peak) of 0.90 ± 0.1 for sustained neurons and 0.66 ± 0.2 for adapting neurons (two-tailed t-test, p = 0.0006; Bonferroni corrected α = 0.00625). These values were measured from current steps that elicited peak hyperpolarization of -90.9 ± 1.4 mV and 91.0 ± 0.4 mV, respectively (Figure 7G_3_). In addition, sustained Y_1_R neurons exhibited a moderate membrane time constant (17.0 ± 8.9 ms), while adapting neurons had a significantly faster membrane time constant (7.3 ± 6.2 ms, Figure 7G_4,_ two-tailed t-test, p = 0.0003; Bonferroni corrected α = 0.00625). Sustained neurons had significantly higher input resistances than adapting neurons (Figure 7G_5_; sustained neurons: R_pk_= 274.2 ± 117.6 MΩ, R_ss_ = 273.2 ± 147.2 MΩ; adapting neurons: R_pk_= 162.8 ± 85.2 MΩ, Rss = 126.3 ± 104.6 MΩ; R_pk_, two-tailed t-test, p = 0.0007; R_ss_, two-tailed t-test, p = 0.0006; Bonferroni corrected α = 0.00625). Y_1_R neurons rarely spontaneously fired at rest and had moderate rheobase values that trended higher for adapting neurons compared to sustained neurons (65.4 ± 50.0 pA for sustained neurons and 95.9 ± 54.2 pA for adapting neurons, Figure 7G_6_, two-tailed t-test, p = 0.0701; Bonferroni corrected α = 0.00625). Based on the numerous significant differences between adapting and sustained Y_1_R neurons, we propose that Y_1_Rs are expressed by at least two distinct types of glutamatergic neurons in the IC.

### NPY hyperpolarizes a subset of Y_1_R-expressing neurons in the IC

NPY signaling is an important modulator of neuronal excitability in several brain regions (Colmers and Bleakman, 1994; Chee et al., 2010; Chen et al., 2019) yet paradoxically during natural behavior these cells are inhibited before feeding begins. Previously, to reconcile these observations, we showed that brief stimulation of AgRP neurons can generate hunger that persists for tens of minutes, but the mechanisms underlying this sustained hunger drive remain unknown (Chen et al., 2016. We therefore hypothesized that NPY signaling alters the excitability of Y_1_R neurons in the IC. To test this hypothesis, we targeted current clamp recordings to Y_1_R neurons while including 1 µM tetrodotoxin in the bath to block endogenous release of NPY from spontaneously active NPY neurons. We found that bath application of [Leu^31^, Pro^34^]-NPY, a high-affinity Y_1_R agonist, hyperpolarized *V*_*rest*_ in 50% of Y_1_R neurons by an average of -6.2 ± 4.3 mV (n = 5 out of 10 neurons; two-tailed paired t-test comparing baseline *V*_*rest*_ to treatment *V*_*rest*_, p = 0.032; Figure 8A). A neuron was considered sensitive to [Leu^31^, Pro^34^]-NPY when *V*_*rest*_ hyperpolarized more than 2 mV below baseline in the presence of the drug. Neurons that did not respond to the drug exhibited a -0.5 ± 0.8 mV change in *V*_*rest*_, which was not significantly different from the baseline *V*_*rest*_ (n= 5 out of 10 neurons; two-tailed paired t-test, p = 0.248). In control experiments performed with a vehicle solution, there was no significant change in *V*_*rest*_ between the baseline and vehicle periods (n = 5, 0.2 ± 1.3 mV; two-tailed paired t-test, p = 0.694; Figure 8B). Neurons that responded to [Leu^31^, Pro^34^]-NPY exhibited a significantly different change in *V*_*rest*_ compared to neurons treated with vehicle (two-tailed Mann-Whitney-Wilcoxon test, p = 0.007; Figure 8C). After a 30 – 50-minute wash-out period, the *V*_*rest*_ of all cells that responded to [Leu^31^, Pro^34^]-NPY returned to baseline (n = 5, two-tailed paired t-test, p = 0.333; data not shown).

We next tested whether faster and more local delivery of [Leu^31^, Pro^34^]-NPY via a puffer pipette would lead to more consistent effects on the *V*_*rest*_ of Y_1_R neurons. Indeed, we found that puff application of [Leu^31^, Pro^34^]-NPY hyperpolarized the *V*_*rest*_ of 89% of Y_1_R neurons by an average of -3.1 ± 2.0 mV (n = 8 out of 9 neurons; two-tailed paired t-test comparing baseline *V*_*rest*_ to treatment *V*_*rest*_, p = 0.007; Figure 8D). In control experiments, there was no significant difference between baseline *V*_*rest*_ and *V*_*rest*_ during vehicle application (n = 7; mean change in *V*_*rest*_ = 0.2 ± 1.1 mV; two-tailed paired t-test comparing baseline *V*_*rest*_ to vehicle *V*_*rest*_, p = 0.57; Figure 8E). As observed in the bath application experiment, there was a significantly different change in *V*_*rest*_ between neurons treated with [Leu^31^, Pro^34^]-NPY and neurons treated with vehicle (two-tailed Mann-Whitney-Wilcoxon test, p = 0.0006; Figure 8F).

Together, these results show that NPY signaling, by hyperpolarizing the resting membrane potential of a population Y_1_R-expressing glutamatergic neurons, provides a novel mechanism to dampen excitability in the IC. Since NPY neurons express NPY (Figure 2 A-C), it is likely that at least some NPY signaling in the IC originates from local NPY neurons.

## Discussion

We identified NPY neurons as a novel class of inhibitory principal neurons in the IC. NPY neurons are GABAergic, produce NPY, are present throughout the IC, and have stellate morphology with dendrites that cross isofrequency laminae in the ICc. NPY neurons account for approximately one-third of GABAergic neurons in the IC and 38 – 50% of stellate neurons in the ICc. NPY neurons fired spontaneously at rest, suggesting they might tonically release GABA and NPY onto postsynaptic targets. Retrograde labeling showed that NPY neurons project to the ipsilateral MG. We also found the first evidence that NPY signaling can regulate excitability in the IC. We showed that Y_1_Rs are expressed by glutamatergic neurons that are widely distributed throughout the IC and which likely consist of two or more neuron types. Intriguingly, most Y_1_R neurons were hyperpolarized by application of a high affinity Y_1_R agonist, suggesting that NPY signaling in the IC might dampen activity through selective effects on an excitatory population of neurons. Thus, NPY neurons represent a major new class of IC inhibitory neurons that may use both GABAergic and NPY signaling to regulate activity levels in the IC and MG.

### NPY neurons are a distinct class of IC neurons

Identification of fundamental neuron classes is essential for understanding the organization and function of neural circuits in the brain. Previous studies showed that the IC contains a large and diverse population of GABAergic neurons. These efforts relied on neuronal morphology (Oliver et al., 1994), in vivo physiology (Ono et al., 2017), neurochemical and extracellular markers (Ito et al., 2009; Beebe et al., 2016) or a combination of morphology, physiology and neurotransmitter content (Ono et al., 2005). However, these studies yielded different results, and it is unclear whether the types of GABAergic neurons they defined represent functionally meaningful groups. Because the neuron groups identified in these studies lack molecular markers, they are not readily accessible for targeted manipulations with optogenetics and other molecular tools.

Combining molecular markers with physiological and anatomical parameters has become key to identifying functionally relevant classes of neurons (Tremblay et al., 2016; Zeng and Sanes, 2017). Through such an approach, we identified NPY neurons as a distinct subtype of GABAergic stellate neurons. NPY neurons are selectively labeled in NPY-hrGFP mice and share a common neurotransmitter content, neuropeptide expression, stellate morphology, and similar physiological features. Ninety-nine percent of NPY neurons immunostained for GAD67, and 95% immunostained for NPY. Ninety-two percent of ICc NPY neurons met the criteria for stellate morphology. In our recordings, 90 – 92% of NPY neurons had a sustained firing pattern. Previously, Ono and colleagues reported that GAD67^+^ neurons exhibited four firing patterns: regular sustained, buildup/pauser, and transient with or without after-depolarizations (Ono et al., 2005). Our results show that NPY neurons fit into the regular sustained category, which included 43% of GAD67^+^ neurons in the Ono study.

Most NPY neurons fired spontaneously at rest. Interestingly, spontaneous firing has been reported for NPY neurons in hypothalamus (Top et al., 2004), striatum (Partridge et al., 2009) and amygdala (Song et al., 2016). This suggests that NPY neurons in multiple brain regions might preferentially express a common suite of ion channels that promotes spontaneous firing. The spontaneous firing of IC NPY neurons indicates that NPY neurons in vivo might provide a constant tone of GABA and NPY to the IC and MG. This inhibitory tone might be rapidly upregulated or downregulated by inputs that modulate the activity levels of NPY neurons, providing a mechanism for NPY neurons to increase or decrease the overall excitability in postsynaptic brain regions.

### NPY neurons include GABAergic projection neurons

In many brain regions, GABAergic neurons are interneurons (Tremblay et al., 2016), but in the IC, GABAergic cells contribute substantially to long-range projections to the MG and other targets (Winer et al., 1996; Peruzzi et al., 1997; Ito et al., 2009; Mellott et al., 2014, 2018; Beebe et al., 2018). The projection to MG is particularly interesting because other regions of primary sensory thalamus do not receive ascending inhibitory input (Halassa and Acsády, 2016). Y_1_R is expressed in the MG (Kishi et al., 2005), and we observed tdTomato expression in a population of MG neurons in Npy1r^cre^ x Ai14 mice (data not shown). Our retrograde experiments confirmed that NPY neurons project to the MG. GABAergic tectothalamic projections arise from multiple IC subdivisions and terminate in multiple MG subdivisions (Mellott et al., 2014), reflecting the parallel organization of the ascending system (Calford and Aitkin, 1983; Rouiller, 1997, Cai et al., 2019). As shown here, NPY has inhibitory effects that could complement GABAergic effects on postsynaptic targets. It will be interesting to determine whether the presence or absence of NPY corelease with GABA underlies some of the differences in collicular-driven inhibition of MG cells (Smith et al., 2007). Since NPY neurons are molecularly identifiable, it is now possible to use optogenetics and other targeted approaches to test how specific inhibitory projections from the IC shape activity in the MG and other targets.

### NPY neurons likely use both GABA and NPY signaling to regulate activity in the IC and MG

NPY is widely expressed in the mammalian brain and is a potent modulator of neuronal activity. Previous brain-wide screening studies suggested that NPY and Y_1_Rs are expressed in the IC (Widdowson, 1993; Kishi et al., 2005; Eva et al., 2006), but the experiments conducted here are the first to characterize IC neurons that synthesize NPY and to test the effects of NPY signaling on IC neurons. Y_1_Rs tend to inhibit neurons when activated (Giesbrecht et al., 2010). Consistent with this, we found that bath application of [Leu^31^, Pro^34^]-NPY hyperpolarized *V*_*rest*_ in 50% of the tested Y_1_ R-expressing neurons, and puff application hyperpolarized *V*_*rest*_ in 89% of the tested neurons. Y_1_R activation can induce opening of G protein-coupled inwardly rectifying potassium (G_IRK_) channels, suggesting that G_IRK_ activation might mediate the effects we observed (Sun et al., 2001; Paredes et al., 2003; Fu et al., 2004; Sosulina et al., 2008), although other mechanisms are possible (Giesbrecht et al., 2010; Villarroel et al., 2018). The fact that not all IC Y_1_R neurons were sensitive to NPY might be due to transient developmental expression of Cre in some neurons. Alternatively, non-responsive neurons might respond to NPY in ways that we did not test.

Beyond influencing auditory computations, the inhibitory effects of NPY signaling might serve diverse roles in the IC. In the hippocampus and amygdala, NPY released from GABAergic NPY neurons acts as an endogenous anticonvulsant (Colmers and Bleakman, 1994; Colmers and El Bahh, 2003; Noe’ et al., 2007). Interestingly, the IC is the main site of initiation of audiogenic seizures (Ross and Coleman, 2000; Faingold, 2002), so NPY/GABA release might protect against hyperexcitability in the IC. Consistent with this, acoustic trauma leads to an increase in the expression of brain-derived neurotrophic factor (BDNF) in the IC (Meltser and Canlon, 2010), and, in other brain regions, BDNF can enhance NPY expression (Nawa et al., 1994; Nobuyuki Takei et al., 1996; Wirth et al., 2005).

The IC also plays an important role in auditory task-related unconditioned fear responses (Muthuraju et al., 2014; Oliveira et al., 2014) and anxiety (Silveira et al., 1993). In the amygdala (Sajdyk et al., 2008), cortex (Vollmer et al., 2016) and hippocampus (Li et al., 2017), NPY exhibits anxiolytic properties and regulates behaviors associated with fear. Thus, NPY neurons in the IC might regulate how IC circuits process unconditioned fear responses and how they are affected by behavioral states like anxiety.

Interestingly, in the IC of Npy1r^cre^ x Ai14 mice we found strong labeling of cells associated with blood vessels (Figure 7A). Previous studies showed that NPY induces contraction in cortical microvessels by activating Y_1_Rs (Abounader et al., 1999; Cauli et al., 2004) and plays a similar role in the cochlea (Carlisle et al., 1990). Since the IC is reported to be the most metabolically active brain region and to have the highest rate of blood flow in the brain (Sokoloff et al., 1977; Gross et al., 1987; Zeller et al., 1997), IC NPY neurons might play a critical role regulating energy homeostasis in the IC.

### NPY and VIP neurons account for more than half of ICc stellate neurons

ICc stellate neurons have long been a subject of speculation, but their functional roles are almost completely unknown due to lack of an approach to target or selectively manipulate them during in vivo recordings or behavioral testing. Here, we have identified a distinct GABAergic cell type labeled in the NPY-hrGFP mouse. In addition, we recently identified VIP neurons as a novel class of glutamatergic stellate cells in the IC, and VIP neurons can be selectively manipulated in VIP-IRES-Cre mice (Goyer et al., 2019). VIP and NPY neurons represent 3.5% and 7.6% of neurons in the ICc, respectively, and combined they account for approximately 55 – 74% of ICc stellate neurons. Thus, for the first time, we are positioned to determine the functional roles of excitatory and inhibitory stellate neurons that together represent the majority of ICc stellate neurons.

## Acknowledgements

We thank Bo Duan for helpful discussions and advice. This work was supported by a Research Grant from the American Hearing Research Foundation (MAS) and National Institutes of Health Grants R56 DC016880 (MTR) and R01 DC004391 (BRS).

